# UBR-1 deficiency leads to ivermectin resistance in *C. elegans*

**DOI:** 10.1101/2024.10.01.616045

**Authors:** Yi Li, Long Gong, Jing Wu, Wesley Hung, Mei Zhen, Shangbang Gao

**Affiliations:** Key Laboratory of Molecular Biophysics of the Ministry of Education, College of Life Science and Technology, Huazhong University of Science and Technology, Wuhan 430074, China; Lunenfeld-Tanenbaum Research Institute, Mount Sinai Hospital, University of Toronto, Toronto, Ontario M5G 1X5, Canada

**Keywords:** Anthelmintic resistance, Ivermectin, UBR-1 deficiency, Glutamate metabolism, GluCls, Ceftriaxone

## Abstract

Resistance to anthelmintics, particularly the macrocyclic lactone ivermectin (IVM), presents a substantial global challenge for parasite control. We found that the functional loss of an evolutionarily conserved E3 ubiquitin ligase, UBR-1, leads to IVM resistance in *Caenorhabditis elegans*. Multiple IVM-inhibiting activities, including viability, body size, pharyngeal pumping, and locomotion, were significantly ameliorated in various *ubr-1* mutants. Interestingly, exogenous application of glutamate induces IVM resistance in wild-type animals. The sensitivity of all IVM-affected phenotypes of *ubr-1* is restored by eliminating proteins associated with glutamate metabolism or signaling: GOT-1, a transaminase that converts aspartate to glutamate, and EAT-4, a vesicular glutamate transporter. We demonstrated that IVM-targeted GluCls (glutamate-gated chloride channels) are downregulated and that the IVM-mediated inhibition of serotonin-activated pharynx Ca^2+^ activity is diminished in *ubr-1*. Additionally, enhancing glutamate uptake in *ubr-1* mutants through ceftriaxone completely restored their IVM sensitivity. Therefore, UBR-1 deficiency-mediated aberrant glutamate signaling leads to ivermectin resistance in *C. elegans*.

## Introduction

Anthelmintic resistance (AR) to broad-spectrum antiparasitic macrocyclic lactones (MLs), which include ivermectin (IVM), avermectin (AVM), and doramectin (DOM), has emerged as a critical global concern in veterinary parasites (*1–3*). The rise of anthelmintic-resistant nematodes poses a pressing issue, impacting not only livestock production but also animal welfare and human health. Overcoming this challenge requires a comprehensive understanding of the molecular mechanisms driving nematode resistance, which, in turn, will facilitate the development of innovative and effective parasite prevention strategies.

The nematode *Caenorhabditis elegans* (*C. elegans*), owing to its sensitivity to a majority of anthelmintic drugs and conservation of functional genes, serves as a crucial model organism for identifying genes related to IVM resistance (*4–7*). Mutations in three *C. elegans* glutamate-gated chloride channel (GluCl) *genes—avr-15*, *avr-14*, and *glc-*1—have collectively demonstrated strong resistance to ivermectin (*8–11*). Exogenous expression and crystallization X-ray structural analysis support the premise that these GluCls act as the primary targets of IVM (*12, 13*). Further identification of IVM resistance-related genes, including P-glycoprotein transporters (*pgp-1*, *pgp-3*, *pgp-6*, *pgp-9*, *pgp-13*, and *mrp-6*), cytochrome oxidases (*cyp-14, cyp-34/35*), and glutathione S-transferases (*gst-4*, *gst-10*) (*14–16*), has revealed their role in enhancing ivermectin excretion or metabolism. These detoxification genes are thought to be regulated by the ubiquitous transcription factor NHR-8, which encodes a nuclear hormone receptor (*17*). While these studies have revealed a polygenic mechanism underlying IVM resistance, increasing reports of unknown causes of IVM resistance continue to emerge (*18–20*), suggesting that previously unrecognized or additional mechanisms regulating GluCls expression may await for further investigation.

UBR1, an E3 ubiquitin ligase and a pivotal component of the ubiquitin-proteasome system, facilitates protein degradation (*21–25*). UBR1 is implicated in substrate-specific recognition and metabolism, primarily through the N-end rule (*26*). In humans, loss-of-function mutations in UBR1 result in Johanson-Blizzard syndrome (JBS), a rare autosomal recessive disorder characterized by a spectrum of developmental and neurological symptoms (*27, 28*). In the nematode *Caenorhabditis elegans*, a single UBR-1 homolog retains all major conserved functional domains (*29*). LIN-28 is the sole identified substrate of *C. elegans* UBR-1, acting as an RNA-binding pluripotency factor crucial for seam cell patterning (*30*). However, the absence of LIN-28 neither replicates nor ameliorates the movement defects observed in *ubr-1* mutants-stiff body bending during backward movement (*29*).

Our recent research has consistently shown that UBR-1 regulates neural outputs through glutamate homeostasis mechanisms. Loss of UBR-1 leads to elevated glutamate levels in animals, disrupting coordinated motor patterns, likely due to a compensatory decrease in the expression of excitatory glutamate receptors (*29*). Furthermore, *ubr-1* mutations disturb inherent asymmetric neural activity by activating inhibitory GluCls (GLC-3 and GLC-2/4), which in turn impairs rhythmic defecation motor programs (*31*). These observations suggest that aberrant glutamate metabolism resulting from UBR-1 deficiency has a widespread effect on glutamatergic signaling pathways. This leads to the question of whether *ubr-1* mutants cause comprehensive changes in IVM-targeted GluCls, thereby contributing to IVM resistance.

To investigate this possibility, we assessed the IVM sensitivity of wild-type and *ubr-1* mutant animals. We found that, in contrast to wild-type N2 animals, loss-of-function *ubr-1* mutants present a range of IVM-resistant phenotypes. N2 animals exhibit high fatality rates, abnormal development and motor defects, such as pharyngeal pumping or locomotion disabilities, upon exposure to ivermectin. In contrast, *ubr-1* mutants were almost insensitive to IVM. The elimination of the glutamate synthetase GOT-1 and the glutamate transporter EAT-4 completely reversed the IVM sensitivity of *ubr-1* mutants, highlighting the critical role of the glutamatergic signaling pathway. Using translational reporters, we found that IVM resistance in *ubr-1* mutants is caused by the functional downregulation of IVM-targeted GluCls, including AVR-15, AVR-14, and GLC-1. These receptors are activated by glutamate to facilitate chloride ion influx into pharyngeal muscle cells, thereby inhibiting muscle contractions and suppressing food intake in *C. elegans*. This downregulation of GluCls is presumably triggered by the aberrantly elevated glutamate-induced compensatory decrease, as the application of exogenous glutamate in wild-type animals partially replicated the IVM resistance observed in *ubr-1* mutants. GluCls downregulation is functionally validated, as evidenced by the diminished IVM-mediated inhibition of serotonin-activated pharyngeal Ca^2+^ activity in *ubr-1* mutants. Moreover, pharmacologically reducing glutamate levels via the use of ceftriaxone (*32, 33*) successfully restored IVM sensitivity in *ubr-1* mutants. Thus, our study presents multiple lines of evidence for a novel IVM resistance mechanism linked to the deficiency of the ubiquitin ligase UBR-1 via a glutamate metabolism pathway.

## Results

### Dose- and time-dependence of IVM resistance in *ubr-1* mutants

To further elucidate the mechanisms of anthelmintic resistance (AR), we employed a simplified assay derived from prior research (**Figure 1A**). We assessed the survival of nematodes on media infused with antiparasitic macrocyclic lactones (MLs). In essence, synchronized L4-stage worms were placed on NGM plates with OP50 bacteria and treated with various MLs, including ivermectin (IVM, 5 ng/mL), avermectin (AVM, 5 ng/mL), or doramectin (DOM, 5 ng/mL). Twenty hours postexposure, we quantified the surviving worms and calculated the percentage of viability relative to the initial population.

**Figure 1.**
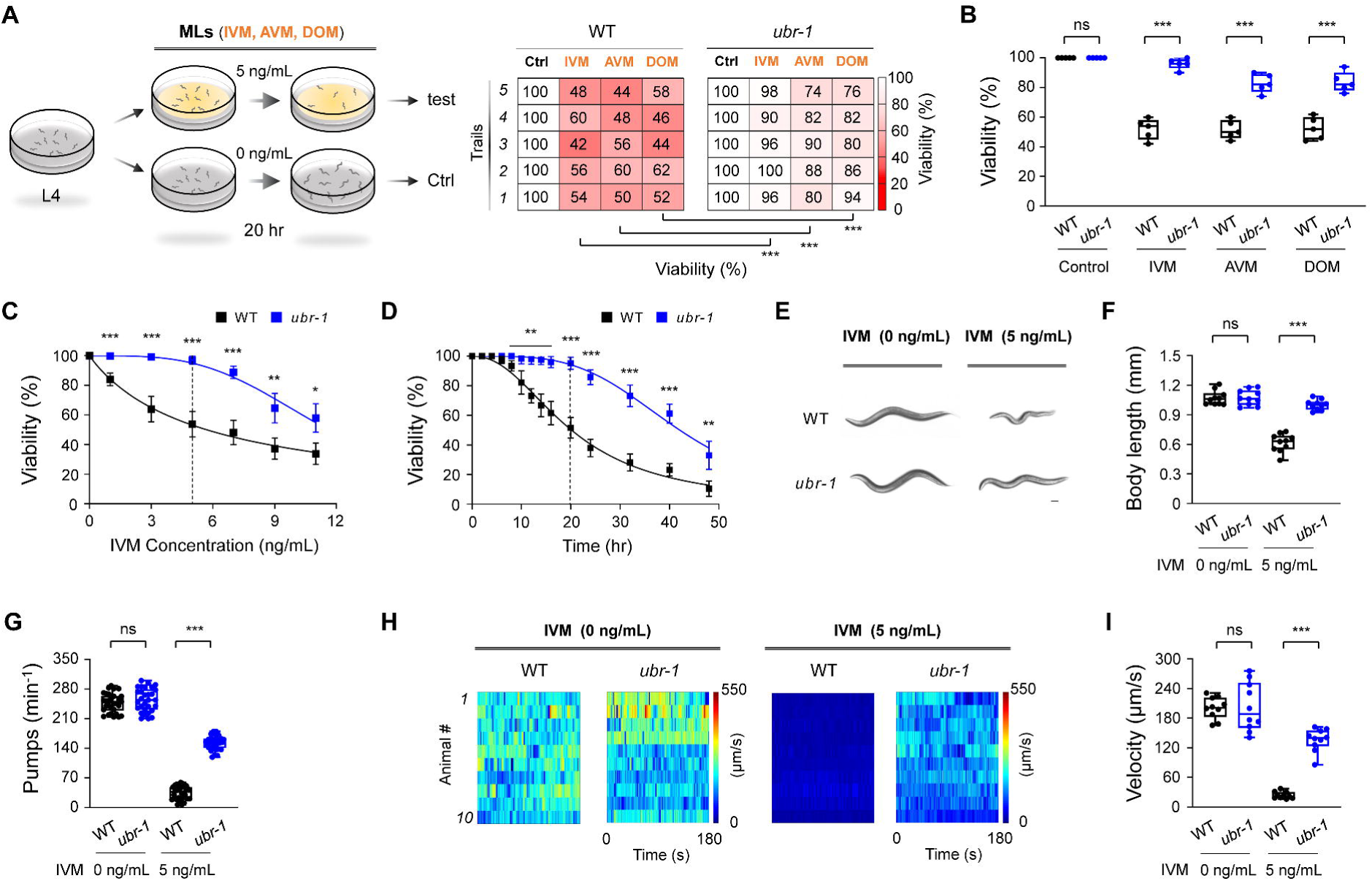
*ubr-1* exhibits IVM-resistant phenotypes. (A) *Left*: Schematic representation of the IVM resistance test in *C. elegans*. Well-fed L4 stage animals were transferred to plates containing OP50 bacteria with or without macrocyclic lactones (MLs) for 20 hours. *Right*: Representative grid plots illustrating the viability of wild-type and *ubr-1* animals in response to ivermectin (IVM, 5 ng/mL), avermectin (AVM, 5 ng/mL), and doramectin (DOM, 5 ng/mL). We used shades of red to represent worm viability on each experimental plate (n = 50 animals per plate), with darker shades indicating lower survival rates. The viability test was repeated at least 5 times (5 trials). (B) Quantification analysis of the viability of wild-type and *ubr-1*(*hp684*) mutants exposed to different MLs. *ubr-1*(*hp684*) mutants exhibit resistance to various MLs. (C) Dose-response curve depicting the viability of wild-type and *ubr-1*(*hp684*) mutants in the presence of varying concentrations of IVM. The IC_50_ was 5.7 ng/mL for the wild type and 11.6 ng/mL for the *ubr-1* mutants. (D) Time-dependent effect of IVM exposure on animal viability. The IC_50_ values were 20.2 hours for the wild type and 42.1 hours for the *ubr-1* mutants. (E) Representative images of worm size in the wild type and *ubr-1*(*hp684*) mutants with or without IVM treatment. Scale bar, 50 µm. (F) Quantitative analysis of body length in different genotypes with or without IVM. (G) Quantification of the average pharynx pump number in animals with or without IVM. (H) Raster plots illustrating the locomotion velocity of individual animals in the absence (0 ng/mL) and presence (5 ng/mL) of IVM. n = 10 animals in each group. (I) Quantification of the average velocity in different genotypes with or without IVM treatment. ns, not significant, *p < 0.05, **p < 0.01, ***p < 0.001 by Student’s *t test*. The error bars represent the SEM.

Our observations revealed a significant decline in the viability of wild-type nematodes following ML exposure (IVM 52.0 ± 3.2%, AVM 51.6 ± 2.9%, DOM 52.4 ± 3.4%, in contrast to the vehicle control 100 ± 0.0%), confirming the susceptibility of *C. elegans* to MLs (**Figure 1B**). In contrast, *ubr-1(hp684; lf)* mutants presented remarkable resistance to ML treatments (IVM, *ubr-1* 96.0 ± 1.7% vs vehicle control 100 ± 0.0%) when juxtaposed with their wild-type counterparts (IVM, WT 52.0 ± 3.2%) (**Figure 1B**). This trend of increased survival was consistently observed with *ubr-1(hp684)* mutants treated with AVM (WT 51.6 ± 2.9% vs *ubr-1* 82.8 ± 2.9%) and DOM (WT 52.4 ± 3.4% vs *ubr-1* 83.6 ± 3.1%). These findings reveal the pronounced ML resistance characteristics of *ubr-1* loss-of-function mutants.

We further investigated the dose- and time-dependent resistance of *ubr-1(hp684)* to select IVM as a proxy for MLs because of its efficiency and common application. We tested a spectrum of IVM concentrations ranging from 1 ng/mL to 11 ng/mL (**Figure S1A**). The data revealed that *ubr-1(hp684)* mutants exhibited resistance at all tested doses (**Figure 1C**), resulting in a doubling of the half-maximal inhibitory concentration (IC_50_) — from 5.7 ng/mL in wild-type N2 worms to 11.6 ng/mL in *ubr-1* mutants. Time-dependent IVM resistance was also compared between the wild type and *ubr-1* mutants (**Figure S1B**). The duration of half-maximal inhibition in the wild type was approximately 20.2 hours, which increased to 42.1 hours in the *ubr-1(hp684)* mutants (**Figure 1D**). To highlight the disparity in IVM resistance, we utilized an assay with 5 ng/mL IVM for a 20-hour period (indicated by the vertical dashed lines in Figure 1C and D) for subsequent experiments, unless otherwise specified.

### Additional IVM resistance phenotypes of *ubr-1*

In addition to viability, *ubr-1* mutants maintained normal growth despite IVM exposure. After 20 hours of IVM, the body size of the wild-type animals significantly decreased (*34*), resulting in a marked decrease in the middle line body length (control, 1.07 ± 0.02 mm; IVM, 0.62 ± 0.03 mm) (**Figure 1E)**. In contrast, the size of the *ubr-1* mutants (control, 1.08 ± 0.02 mm; IVM, 1.00 ± 0.02 mm) was largely preserved, indicating that the *ubr-1* mutants developed into typical young adults (**Figure 1F)**. Notably, while IVM exposure nearly halted pharyngeal pumping in wild-type animals (control 247.4 ± 4.1 min^-1^, IVM 34.6 ± 2.8 min^-1^), *ubr-1* mutants sustained considerable pumping activity (control 254.1 ± 5.0 min^-1^, IVM 152.9 ± 3.0 min^-1^) (**Figure 1G**). Motor functions were similarly resistant to IVM in *ubr-1* mutants (**Movie S1**). The velocity of free movement under IVM treatment was significantly greater in *ubr-1* mutants (135.9 ± 7.2 μm/s) than in wild-type animals (23.8 ± 2.1 μm/s) (**Figure 1H, I, Figure S1C**). No significant differences in viability, body length, pharyngeal pumping, or locomotion speed were detected between *ubr-1* and wild-type animals in the absence of IVM treatment.

These results collectively indicate that *ubr-1* loss-of-function mutants present a comprehensive spectrum of phenotypes associated with IVM resistance.

### Different *ubr-1* mutants exhibit consistent IVM resistance

The *ubr-1* gene is functionally conserved from yeast to humans. In nematodes or filarial worms, *ubr-1* also exhibited high sequence similarity (**Figure S2A**). The *hp684* allele contains a premature stop codon, resulting in the truncation of the final 194 amino acids of UBR-1 (Q1864X) (**Figure S2B**). To validate functional congruence, we expanded our study to include additional *ubr-1* mutant alleles. These alleles, which are genetically predicted to lack one or more critical conserved domains, including *hp865*, which replaces the entire RING finger domain with an SL2-NLS::GFP, and *hp821 hp833* (E34X, E1315X), each introducing stop codons that suggest the absence of all functional domains. All these alleles demonstrated IVM resistance comparable to that of the *hp684* allele. Specifically, the viability of *hp865* (81.6 ± 3.8%) and *hp821 hp833* (76.4 ± 2.6%) increased following IVM treatment (**Figure S2C**). Similarly, the body lengths of *hp865* (0.93 ± 0.02 mm) and *hp821 hp833* (0.90 ± 0.02 mm) were not diminished by IVM exposure (**Figure S2D**). Pharyngeal pumping and spontaneous movement velocities were also significantly greater in *hp865* (154.2 ± 3.4 min^-1^, 100.2 ± 7.4 μm/s) and *hp821 hp833* (134.9 ± 3.7 min^-1^, 86.1 ± 6.9 μm/s) than in the wild type (**Figure S2E, F**). These findings demonstrate that various *ubr-1* mutants exhibit a consistent IVM-resistant phenotype. While we have identified relevant sequences in parasitic nematodes including *Onchocerca volvulus*, *Brugia malayi*, and *Toxocara canis*, potential mutations in *ubr-1*-like genes in these parasitic nematodes may lead to ivermectin resistance.

### Glutamate induces IVM resistance in wild-type worms

Systemic elevation of glutamate levels in *ubr-1* mutants has been documented (*29, 35, 36*), prompting an investigation into whether this increase is the key determinant of the observed resistance to IVM.

To test this hypothesis, we pretreated wild-type N2 animals with glutamate to mimic the excessive glutamate environment in *ubr-1* mutants (**Figure 2A**). During these trials, wild-type larvae were exposed to L-glutamate (20 mM) on NGM plates from the L1 to L4 stages before their transfer to IVM-containing plates. Strikingly, these glutamate-pretreated animals presented a significant increase in IVM resistance (**Figure 2B**).

**Figure 2.**
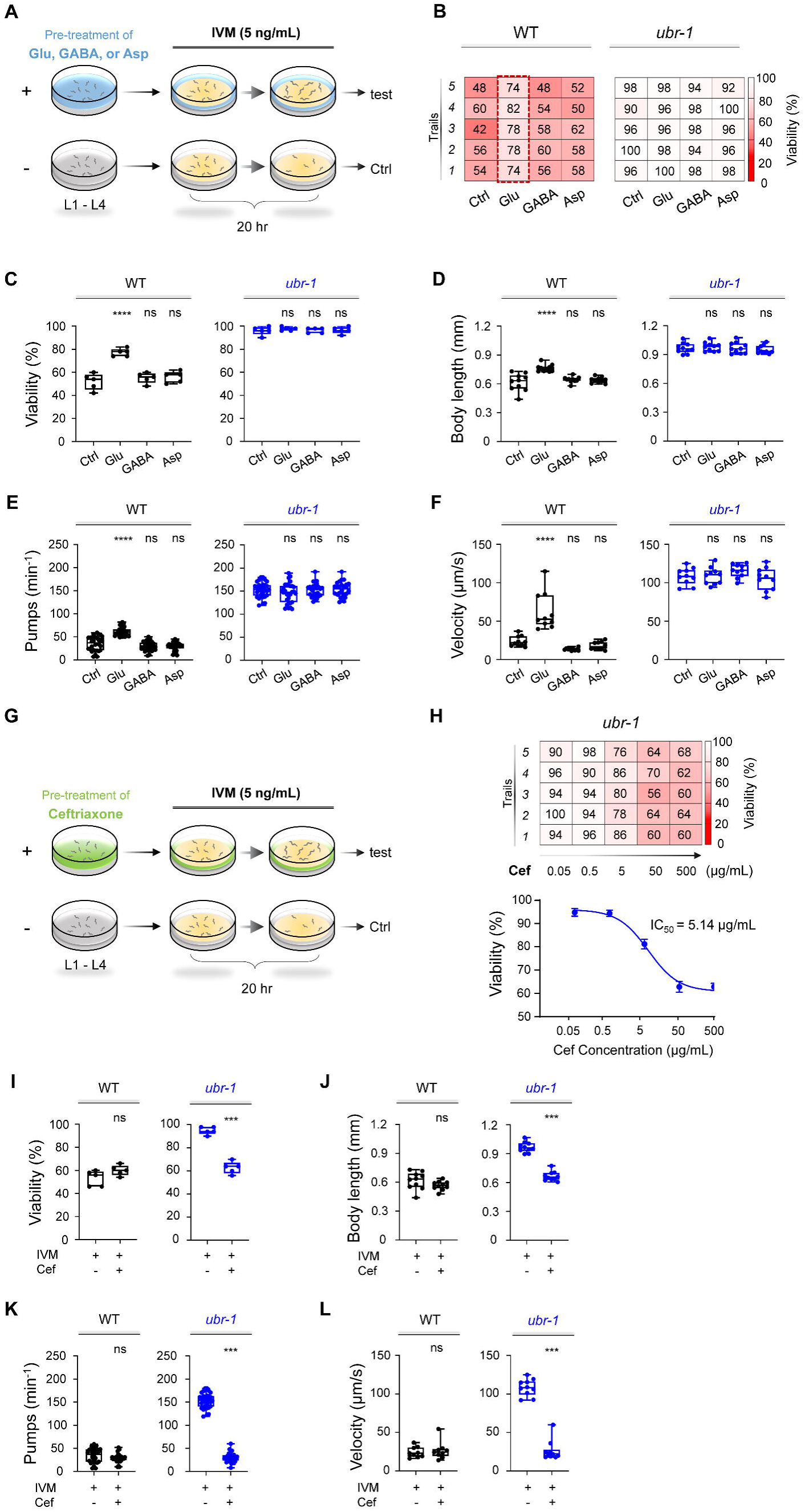
Glutamate mimics IVM resistance in wild-type N2 animals, and ceftriaxone fully reverses IVM resistance in *ubr-1* mutants. (A) Modified schematic representation of the pretreatment approach in the IVM resistance test in *C. elegans*. Synchronized L1 animals were cultured on plates containing OP50 bacteria and pretreated with glutamate (Glu, 5 mM), γ-aminobutyric acid (GABA, 5 mM), or aspartate (Asp, 5 mM) until they reached the L4 stage. Pretreated animals were then transferred to plates containing additional IVM (5 ng/mL) for 20 hours. (B) Representative grid plots illustrating the viability of wild-type and *ubr-1* animals pretreated with different glutamate metabolites. (C-F) Quantitative analysis of viability, body length, pharyngeal pump rate, and locomotion velocity in wild-type and *ubr-1* mutants following exposure to various glutamate metabolites. Wild-type worms treated with glutamate displayed resistance to IVM. However, glutamate did not affect the IVM resistance of the *ubr-1* mutant. ns, not significant, ****p < 0.0001 by one-way ANOVA in C-F. (G) Diagram illustrating the ceftriaxone pretreatment procedure in the IVM resistance test. Synchronized L1 animals were cultured on OP50-fed plates with or without ceftriaxone (50 μg/mL) until the L4 stage. The animals were subsequently transferred to plates seeded with additional IVM (5 ng/mL) for 20 hours. (H) *Upper*: Representative grid plots depicting the viability of *ubr-1* animals at different concentrations of ceftriaxone. *Bottom*: Dose-response curve illustrating the amount of ceftriaxone required for viability in *ubr-1*(*hp684*) mutants (IC_50_ = 5.1 μg/mL). (I-L) Quantification of viability, body length, pharyngeal pumping, and locomotion velocity in the wild type and *ubr-1* mutants in the absence and presence of ceftriaxone. Ceftriaxone fully restored the sensitivity of *ubr-1* mutants to IVM, comparable to the levels observed in wild-type N2 animals. ns, not significant, ***p < 0.001 by Student’s *t test* in I-L. Error bars represent SEM.

Specifically, the viability of these glutamate-exposed animals increased from 52 ± 3.2% to 77.2 ± 1.5% compared with that of their nonexposed counterparts (**Figure 2C**). A corresponding increase in body size was noted, from 0.62 ± 0.03 mm to 0.76 ± 0.01 mm (**Figure 2D**). Motor activity similarly improved, with velocities increasing from 23.79 ± 2.1 μm/s to 62.4 ± 7.6 μm/s (**Figure 2E-F**). These findings clearly demonstrate that the external application of glutamate can induce IVM resistance in wild-type N2 animals. Interestingly, in *ubr-1* mutants, pretreatment with glutamate did not further increase IVM resistance, implying potential saturation of the glutamate effect in *ubr-1* mutants (**Figure 2B-F**).

In addition to glutamate, other metabolically related molecules, such as aspartate (Asp), a precursor in glutamate synthesis, and γ-aminobutyric acid (GABA), a product of glutamate metabolism, were also found to be elevated in the *ubr-1* mutant (*29, 31*). However, the cocultivation of wild-type N2 worms with aspartate or GABA did not confer IVM resistance (**Figure 2A-F**), suggesting the glutamate-specific regulation of IVM resistance. This finding highlights the specificity and sufficiency of glutamate in conferring IVM resistance in a wild-type context.

### Enhancing glutamate uptake restores IVM sensitivity in *ubr-1*

Given the likelihood that elevated glutamate levels underpin IVM resistance in *ubr-1*, mitigating this glutamate surplus could reverse resistance and reinstate IVM sensitivity. To evaluate this premise, we employed ceftriaxone (Cef) (**Figure 2G)**, which is known for enhancing glutamate uptake by upregulating excitatory amino acid transporter-2 (EAAT2) (*32, 33*), with the aim of reducing glutamate levels.

The results were striking: the IVM resistance typically observed in *ubr-1* mutants was completely counteracted by Cef treatment. While Cef (50 μg/mL) had a negligible effect on wild-type N2 animals, it fully restored IVM sensitivity in *ubr-1* mutants (**Figure 2H**). Moreover, this restoration followed a dose-dependent pattern, with a half-effect concentration of 5.14 μg/mL (**Figure 2H**). Cef treatment resulted in a decrease in the viability of *ubr-1* mutants from 96.0 ± 1.7% to 62.8 ± 2.3% in the presence of IVM (**Figure 2I**). Additional IVM resistance phenotypes in *ubr-1* mutants, such as body size, pharyngeal pumping frequency, and locomotion velocity, were fully ameliorated by Cef (**Figure 2J-L**).

In line with its classification as a beta-lactam antibiotic, Cef effectively reduced the growth of the worm food source *E. coli* OP50, resulting in a reduced bacterial lawn on NGM plates (**Figure S3A, B**). However, simply diminishing the OP50 bacterial quantity on NGM plates without Cef did not restore IVM sensitivity in *ubr-1* mutants (**Figure S3C-F**), suggesting that food scarcity is not a critical factor. Similarly, increasing the number of OP50 bacteria present on NGM plates containing Cef did not affect its ability to restore IVM sensitivity in *ubr-1* mutants (**Figure S3C-F**). These findings indicate that the antimicrobial action of ceftriaxone is not related to its ability to restore IVM sensitivity in *ubr-1* mutants.

Hence, these results indicate that pharmacologically reducing glutamate levels in *ubr-1* reestablishes IVM sensitivity, leading to the hypothesis that aberrant glutamate metabolism underlies IVM resistance in *ubr-1*.

### Restoration of *ubr-*1 IVM sensitivity through glutamatergic signaling pathway regulation

Our previous research revealed the involvement of UBR-1 in regulating glutamate metabolism, which is essential for coordinated reversal bending and rhythmic defecation motor programs (*29*). The absence of UBR-1 also leads to perturbations in the balance between the GABAergic and glutamatergic signaling pathways (*31*). In *C. elegans*, IVM sensitivity is marked by the sustained activation of glutamate-gated chloride channels (GluCls) by IVM (*11*). These glutamate-related findings led us to explore whether the resistance of *ubr-1* mutants to IVM could be attributed to alterations in the glutamate pathway.

At glutamatergic synapses, a series of enzymatic reactions link glutamate to neurotransmitter recycling. In *C. elegans*, glutamate-oxaloacetate transaminases, or GOT enzymes, facilitate amino group conversion between aspartate and α-ketoglutarate (α-KG) to oxaloacetate (OAA) and glutamate (**Figure 3A**) (*37, 38*). We began our exploration by assessing IVM sensitivity following the disruption of GOT-1. While single *got-1* loss-of-function mutants displayed IVM sensitivity similar to that of the wild type, *ubr-1; got-1* double mutants presented a significant increase in IVM sensitivity, reverting to wild-type levels of viability, body length, and motor activity, in stark contrast to *ubr-1* single mutants (**Figure 3B-E**). These findings indicate that impeding glutamate synthesis can effectively restore IVM sensitivity in *ubr-1* mutants.

**Figure 3.**
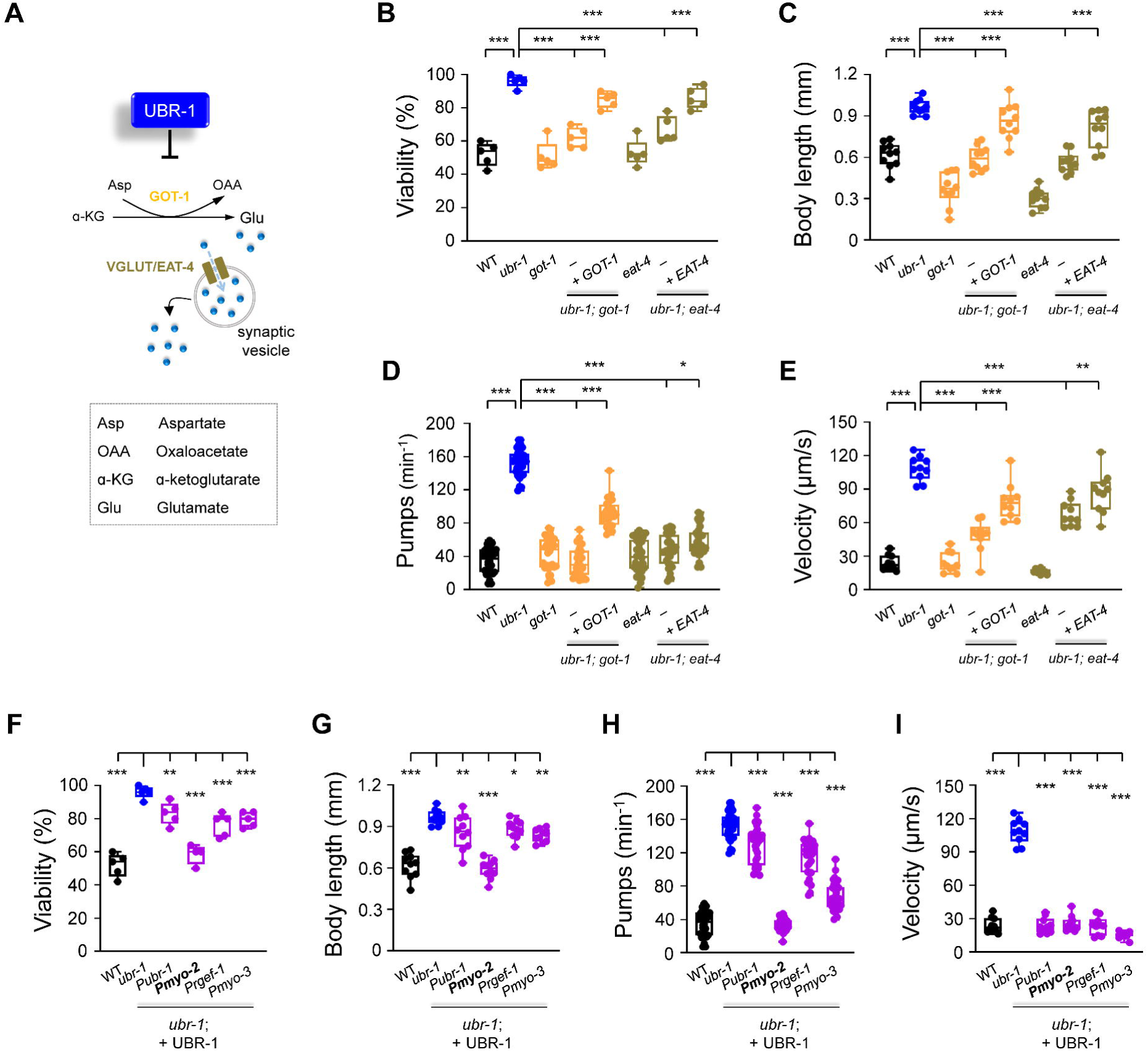
Deletion of *got-1* and *eat-4* suppresses *ubr-1*’s IVM resistance. (A) Schematic of the pathway for glutamate synthesis through transaminase GOT-1 and loading via vesicular transport VGLUT/EAT-4. (B-E) Quantification of viability, body length, pharyngeal pump rate, and locomotion velocity in different genotypes. The removal of *got-1* and *eat-4* significantly restored sensitivity to IVM (5 ng/mL) in *ubr-1* mutants, whereas the re-expression of GOT-1 (+GOT-1) and EAT-4 (+EAT-4) partially reinstated IVM resistance in the respective double mutants. (F-I) IVM resistance phenotypes were rescued by expression of UBR-1 (+UBR-1) driven by its own promoter (P*ubr-1*) or the pharynx muscle-specific promoter (P*myo-2*). For the viability test in F, n = 50 animals per plate and repeated at least 5 times (5 trials), n ≥ 10 animals in G-I, *p < 0.05, **p < 0.01, ***p < 0.001 by one-way ANOVA. All statistical analyses were performed against *ubr-1* mutant. Error bars, SEM.

Glutamate, a crucial excitatory neurotransmitter, is packed into synaptic vesicles by the vesicular glutamate transporter (VGLUT) and released into the synaptic cleft to activate postsynaptic receptors. We next explored the impact of inhibiting glutamate transport in *ubr-1* mutants by eliminating EAT-4, the sole VGLUT in *C. elegans* (*39*). Notably, the absence of EAT-4 in *ubr-1* mutants dramatically reversed their IVM resistance, as evidenced by increased viability (*ubr-1; eat-4* 66.8 ± 3.5%), reduced body length (*ubr-1; eat-4* 0.56 ± 0.02 mm), increased pharyngeal pumping (*ubr-1; eat-4* 46.7 ± 3.6 min^-1^), and improved locomotion speed (*ubr-1; eat-4* 83.5 ± 7.4 μm/s) in *ubr-1; eat-4* double mutants (**Figure 3B-E**).

In summary, these results underscore the importance of glutamate synthesis and synaptic transport in restoring IVM sensitivity in *ubr-1* mutants, suggesting a role for synaptic glutamate signaling in the resistance of *ubr-1* to IVM. The IVM resistance phenotypes of wild-type N2 animals were not rescued by ceftriaxone (**Figure 2I-L**), suggesting that ceftriaxone appears to be effective only against excess glutamate. This discovery is particularly intriguing, as it reveals a previously unexplored role of glutamate metabolism in the regulation of IVM resistance.

### UBR-1 appears to regulate IVM resistance from the pharynx

UBR-1 is ubiquitously expressed across a variety of tissues, including the pharynx, body wall muscles, and neurons (**Figure S4A**) (*29, 35, 36*). To pinpoint the critical tissue for the role of UBR-1 in IVM resistance, we conducted tissue-specific rescue assays via a functional plasmid. Our findings revealed that restoring UBR-1 expression under its endogenous promoter attenuated the IVM resistance observed in *ubr-1* mutants. Notably, when UBR-1 expression was driven by an exogenous pharyngeal muscle-specific promoter, it fully corrected the IVM resistance phenotypes of the *ubr-1* mutants (**Figure 3F-I**). These results imply that although UBR-1 is important in multiple tissues, its role in the pharynx is particularly critical in modulating IVM sensitivity.

To verify the tissue-specific function of UBR-1, we utilized RNA interference (RNAi) to selectively decrease *ubr-1* expression in the pharynx. This precise RNAi application to the pharynx indeed induced IVM resistance in wild-type N2 worms (**Figure S4B-E**). RNAi-mediated knockdown of *ubr-1* in nonpharyngeal tissues, such as neurons and body wall muscles, also elicits an IVM resistance profile. Our research revealed that animals in which *ubr-1* was knocked down specifically in the pharynx presented more pronounced IVM resistance phenotypes than did those in which *ubr-1* was knocked down in other tissues. This phenotype approached the level observed when *ubr-1* was knocked down via its native promoter, underscoring the unique contribution of the pharynx to *ubr-1*-mediated IVM resistance.

Taken together, these results confirm that UBR-1 predominantly regulates IVM resistance through its action in the pharynx.

### Downregulation of IVM-targeted GluCls in *ubr-1*

What mechanisms underlie the induction of IVM resistance by abnormal glutamate metabolism? Previously, we established that *ubr-1* mutants exhibit impaired GLR-1 glutamate receptor function in AVA/AVE interneurons (*29*). Three GluCls (*avr-15*, *avr-14*, *glc-1*) are recognized as pivotal for IVM resistance in *C. elegans* (*11*). We sought to determine whether compromised IVM-targeted GluCls contribute to IVM resistance in *ubr-1* mutants.

To test this hypothesis, we created three transgenic strains with translational reporters for each GluCl under the control of their native promoters (P*avr-15*::AVR-15::GFP, P*avr-14*::AVR-14::GFP, P*glc-1*::GLC-1::GFP). Notably, AVR-15::GFP expression was pronounced in the pharynx, whereas GLC-1::GFP displayed a distinct head expression pattern, particularly concentrated in the head body wall muscles (**Figure 4A, Figure S5A**). AVR-14::GFP expression was localized primarily to various head and body neurons (**Figure S5A**). Intriguingly, compared with that in wild-type animals, the fluorescence intensity of these GluCls was markedly lower in *ubr-1* mutants (**Figure S5A, B**). The modulation of GluCl expression by *ubr-1* seems to be posttranscriptional, given that the RNA levels of the GluCl-encoding genes (*avr-15*, *avr-14*, *glc-1*) did not significantly differ (**Figure S5C**).

**Figure 4.**
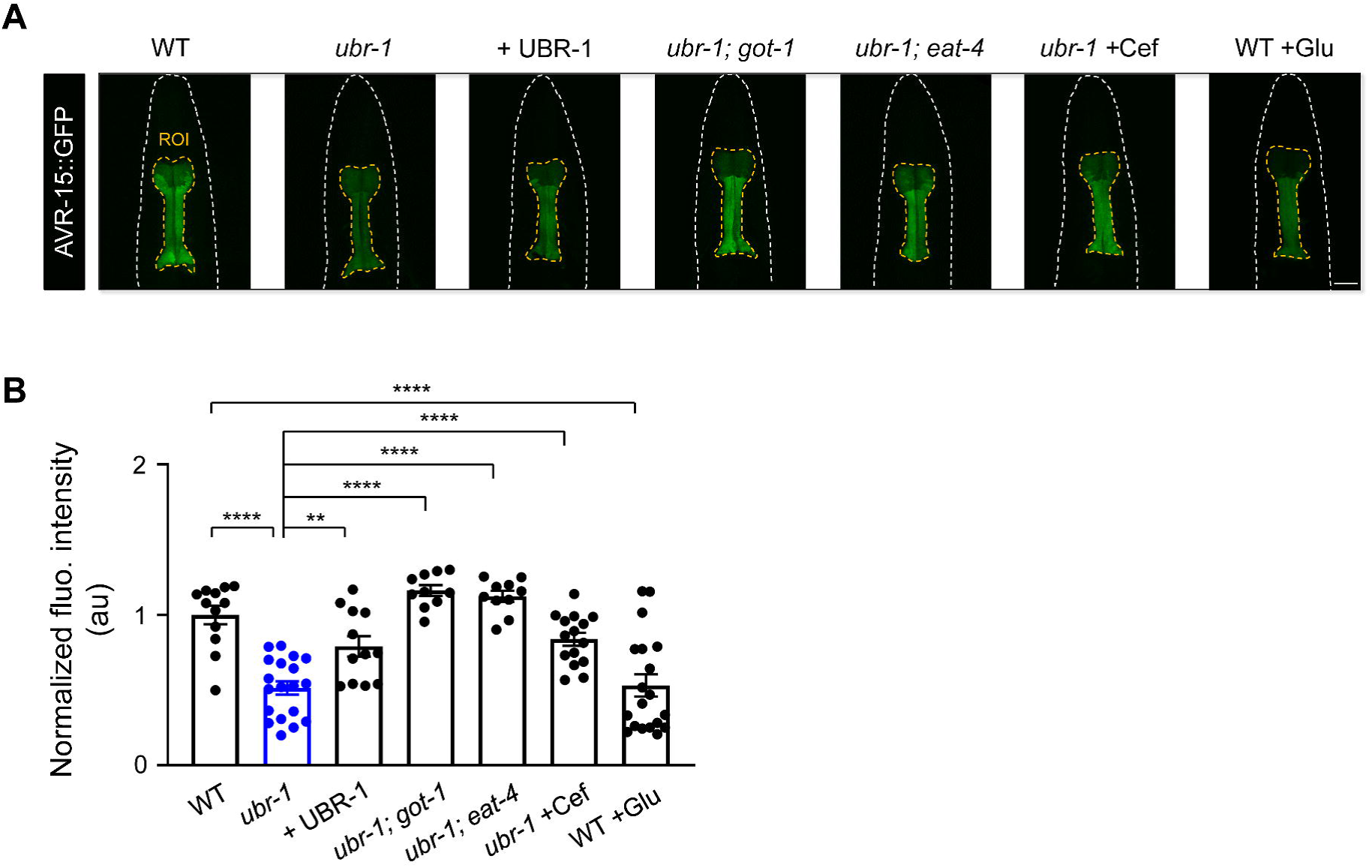
Downregulation of AVR-15 in *ubr-1* mutants. (A) Representative fluorescence intensity of AVR-15 in different genotypes. (B) Quantitative analysis of normalized fluorescence intensity revealed a reduction in AVR-15::GFP expression in *ubr-1* mutants. The diminished AVR-15 levels were rescued by overexpression of UBR-1 and genetically suppressed in *got-1* and *eat-4* mutants. Pharmacological intervention with ceftriaxone (50 μg/mL) successfully reinstated the pharyngeal expression of AVR-15::GFP. Conversely, glutamate administration reduced AVR-15 expression in the wild-type N2 strain. ROI, region of interest; n ≥ 10 animals in each group. **p < 0.01, ****p < 0.0001 by one-way ANOVA. Error bars, SEM.

Crucially, the reduction in pharyngeal AVR-15::GFP fluorescence observed in *ubr-1* mutants was reversed upon reintroduction of UBR-1 (**Figure 4A**). Additionally, eliminating *got-1* and *eat-4* in these mutants mitigated the decrease in AVR-15::GFP expression within the pharynx (**Figure 4A, B**). Overexpression of AVR-15 also partially restored the sensitivity of ubr-1 mutant to IVM (**Figure S5D**). These results suggest a direct role of glutamate signaling in regulating GluCl expression levels. Notably, ceftriaxone pretreatment effectively reversed the diminished pharyngeal AVR-15::GFP levels in *ubr-1* mutants (**Figure 4A, B**). In addition to glutamate pretreatment eliciting IVM resistance in wild-type animals, the application of glutamate markedly diminished the pharyngeal AVR-15::GFP fluorescence in these same wild-type worms (**Figure 4A, B**). Interestingly, the ability of ceftriaxone pretreatment to rescue viability in *ubr-1* mutants was completely lost in the *avr-14; avr-15; glc-1* triple mutants (**Figure S5E**), confirming that ceftriaxone restores IVM sensitivity by reducing glutamate levels, which in turn upregulates GluCl receptors.

To further investigate the potential regulatory influence of *ubr-1* on a broader spectrum of synaptic receptors, we selected two distinct categories of receptors that are not activated by glutamate. This reduction was not observed for the excitatory acetylcholine receptor (UNC-29/AChR) or inhibitory GABA receptor (UNC-49/GABA_A_R) (**Figure S6**), suggesting that *ubr-1* may selectively affect glutamate-gated receptors (*29*).

Collectively, these results demonstrate that aberrant glutamate metabolism in the *ubr-1* mutant leads to the downregulation of IVM-targeted GluCls, which likely contributes to diminished IVM responsiveness, culminating in the IVM resistance observed in *ubr-1* mutants.

### IVM completely suppresses serotonin-evoked pharynx activity in wild-type animals but not in *ubr-1*

IVM acts by engaging target GluCls, leading to the irreversible inhibition of pharyngeal pumping, disrupting conduction between nerve and muscle cells and eventually killing parasites (*8, 11*). The intricate dynamics of pharyngeal pumping in *C. elegans* rely on the coordinated interaction between the pharyngeal muscles and the excitatory pacemaker neuron MC, which is activated by serotonin (*40*).

To further examine the functional impairments of IVM-targeted GluCls, we conducted a direct examination of the impact of IVM on pharyngeal muscle cell activity. We generated a transgenic line expressing a Ca^2+^ sensor, GCaMP6, in the pharynx, enabling real-time monitoring of pharyngeal muscle activity (**Figure 5A, Methods**). Using 5-HT to stimulate excitatory MC neurons, we induced intrinsic Ca^2+^ activity within the pharyngeal muscles. In these assays, 5-HT triggered strong Ca^2+^ transients across the pharyngeal muscle cells, from the terminal bulb through the isthmus to the anterior bulb (**Figure 5A**). To align with our quantified pumping behavior analysis, we focused our observations on the Ca^2+^ intensity within the terminal bulb. We noted a dose-dependent response of pharyngeal muscle cells to 5-HT, with an estimated EC_50_ of approximately 20.1 mM (**Figure 5B, C**). This assay provides a practical method to evaluate the functional impairment of IVM-targeted GluCls in *ubr-1*.

**Figure 5.**
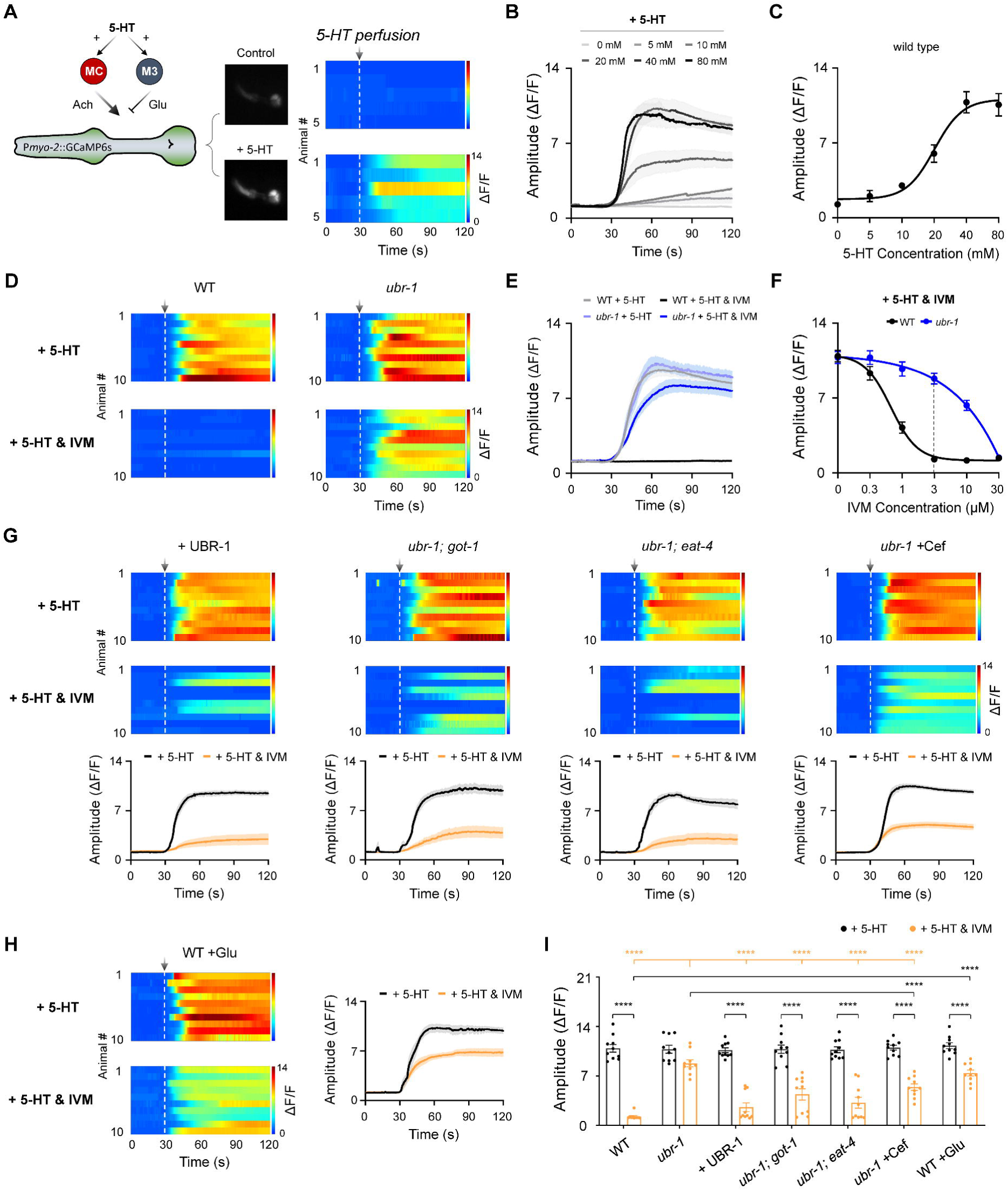
Reduced inhibition of serotonin-stimulated pharyngeal activity by IVM in *ubr-1* mutants. (A) *Left*: Schematic diagram of the pharynx motor circuit composed of an excitatory cholinergic motor neuron MC, an inhibitory glutamatergic motor neuron M3 and a group of compact muscles. Both types of motor neurons could be activated by serotonin (5-HT). *Right*: Representative pharynx muscle Ca^2+^ response and kymograph evoked by 5-HT (20 mM). The dashed lines aligned with the arrows denote the initiation of perfusion. (B) Average Ca^2+^ traces evoked by different concentrations of 5-HT. (C) Dose-response curve of 5-HT for the pharynx muscle Ca^2+^ response in wild-type animals (EC_50_ = 20.1 mM). n ≥ 5 animals in each group. (D) Baseline-subtracted kymograph of the 5-HT-evoked pharynx muscle Ca^2+^ response in wild-type and *ubr-1* mutants. IVM (3 μM) completely inhibited the wild-type pharynx Ca^2+^ response, which was significantly reduced in *ubr-1* mutants. (E) Average Ca^2+^ traces evoked by 5-HT in different genotypes with or without IVM. (F) The dose-dependent inhibitory effect of IVM on pharynx Ca^2+^ was attenuated in *ubr-1* mutants (blue line) compared with wild-type animals (black line) (IC_50_ = 0.5 μM in WT, while IC_50_ = 10.1 μM in *ubr-1*). (G) The decreased IVM inhibition of serotonin-evoked pharynx activity in *ubr-1* was rescued by the restoration of UBR-1 expression (+ UBR-1), genetic removal of *got-1* and *eat-4*, and pretreatment with ceftriaxone (+ Cef, 50 μg/mL). (H) IVM-mediated inhibition of serotonin-evoked pharynx activity in wild-type animals was blocked by glutamate feeding (WT + Glu, 20 mM). n = 10 animals in each group. (I) Quantification of the 5-HT-evoked peak amplitude of the pharynx muscle Ca^2+^ response in the different genotypes. Black stars: ****p < 0.0001 was used to compare intragroup differences or pharmacological groups via Student’s *t test*; apricot stars: ****p < 0.0001 was used to compare the differences with *ubr-1* in the “+ 5-HT & IVM” group via one-way ANOVA. Error bars, SEM.

Interestingly, we observed that 5-HT induced similar Ca^2+^ transients in both wild-type and *ubr-1* mutant worms. Furthermore, upon application of IVM (3 μM), 5-HT-elicited Ca^2+^ activity was profoundly suppressed in the wild-type pharynx (**Figure 5D**), mirroring its effect on pharyngeal pumping. However, *ubr-1* mutants exhibited marked resistance to the suppressive effects of IVM, with only a modest decrease in the amplitude of Ca^2+^ transients (**Figure 5E**). Essentially, the capacity of IVM to inhibit 5-HT-driven pharynx activity was significantly compromised in *ubr-1* mutants. Additionally, consistent with its behavioral impact, IVM dose-dependently suppressed 5-HT-elicited pharynx Ca^2+^ activity, with an estimated IVM IC_50_ of approximately 0.5 μM for wild-type and 10.1 μM for *ubr-1* mutants (**Figure 5F**). These findings reveal the functional deficits of IVM-targeted GluCls in *ubr-1*.

In line with our earlier behavioral findings, the restoration of UBR-1 expression successfully reversed the inhibition of 5-HT-induced pharynx Ca^2+^ by IVM in *ubr-1* mutants (**Figure 5G**). This corrective effect was similarly observed in *ubr-1; got-1* and *ubr-1; eat-4* double mutants (**Figure 5G, I**), reinforcing the association between decreased GluCl expression and attenuated IVM inhibition in *ubr-1* mutants. These results suggest a mechanism whereby elevated glutamate levels in *ubr-1* mutants may lead to glutamate-induced downregulation of GluCls, ultimately leading to IVM resistance.

To substantiate this proposed mechanism, we evaluated pharyngeal Ca^2+^ activity in *ubr-1* mutants pretreated with Cef and in wild-type animals pretreated with glutamate. Consistent with our behavioral data, Cef treatment effectively reversed the inhibitory effect of IVM on 5-HT-evoked Ca^2+^ activity in the pharynx of *ubr-1* mutants (**Figure 5G, I**), whereas glutamate exposure resulted in significant IVM resistance in wild-type animals (**Figure 5H, I**). This alignment between behavioral outcomes and Ca^2+^ activity measurements further supports the notion that glutamate accumulation is a critical factor in the downregulation of GluCls and the ensuing development of IVM resistance in *ubr-1* mutants.

## Discussion

In this study, we identified a crucial role of the E3 ubiquitin ligase UBR-1 in *C. elegans*, which contributes to IVM resistance. Given the importance of UBR-1 in regulating glutamate metabolism, our research introduces a novel mechanism for IVM resistance that could be applicable to other macrocyclic lactone (ML) anthelmintic resistance scenarios. Notably, this glutamate-based mechanism operates upstream, affecting the expression levels of GluCls, a departure from the previously documented variations in detoxification transporters that facilitate drug efflux. Through detailed investigation, we demonstrated that UBR-1 is instrumental in maintaining physiological glutamate homeostasis, thereby governing the abundance of various glutamate receptors. In *ubr-1* mutants, perturbations in glutamate metabolism lead to reduced expression of primary IVM-sensitive GluCls, leading to decreased IVM-induced inhibition of serotonin (5-HT)-evoked pharynx Ca^2+^ activity and subsequent resistance to IVM.

### A working model for UBR-1-mediated IVM resistance in *C. elegans*

Building upon our findings, we propose a working model that illustrates the role of the E3 ubiquitin ligase UBR-1 in mediating IVM resistance (**Figure 6**). In *C. elegans*, UBR-1 ensures functional glutamate homeostasis by inhibiting the activity of transaminase GOT-1 through a pathway that has yet to be resolved. A deficiency in UBR-1 leads to uncontrolled production of excess glutamate by GOT-1, which is then packed into synaptic vesicles for release via the vesicular glutamate transporter VGLUT/EAT-4. This excess glutamate results in the downregulation of glutamate receptors, including the IVM-targeted GluCls AVR-15, AVR-14, and GLC-1. The consequent reduction in these receptors diminishes the inhibitory effect of IVM on 5-HT-evoked pharynx Ca^2+^ activity, highlighting the notable IVM resistance in *ubr-1* mutants. Furthermore, the exogenous application of glutamate induces IVM resistance in wild-type animals, whereas the pharmacological enhancement of glutamate transporter activity via ceftriaxone mitigates resistance in *ubr-1* mutants. These observations support the conclusion that IVM resistance in *ubr-1* mutants is intricately linked with aberrant glutamate metabolism.

**Figure 6.**
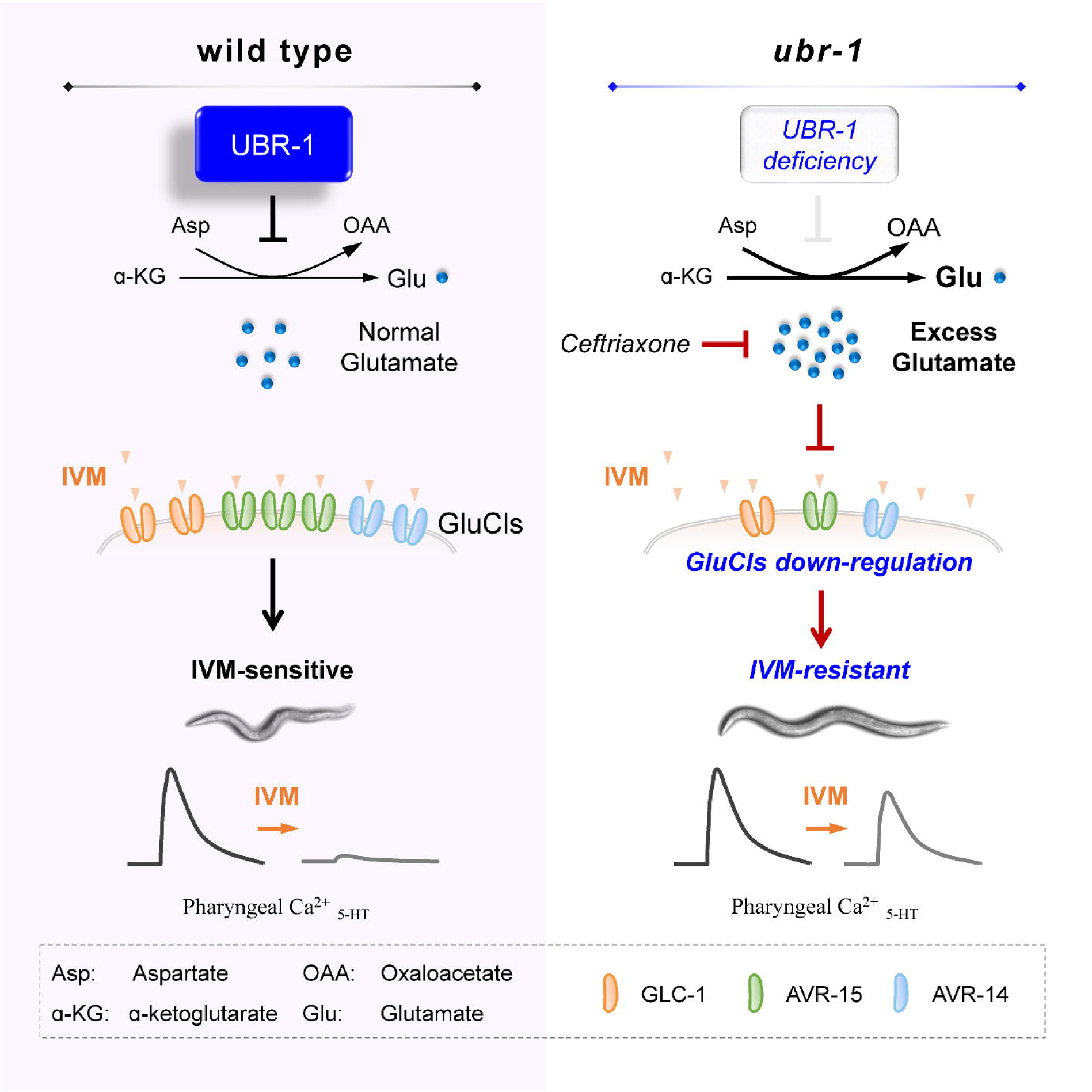
A working model. In wild-type animals, functional UBR-1 helps maintain balanced glutamate levels by inhibiting transaminase GOT-1 activity. In contrast, in *ubr-1* loss-of-function mutants, the absence of GOT-1 inhibition leads to excessive glutamate production. This glutamate excess, which can be reduced by ceftriaxone treatment, induces compensatory downregulation of IVM-targeted GluCls. This downregulation results in a diminished inhibitory response to IVM in the pharyngeal region, ultimately causing IVM resistance in *ubr-1* mutants.

### Unveiling a novel mechanism of anthelmintic resistance: aberrant glutamate metabolism

Glutamate is a central metabolic hub that connects glucose and amino acid metabolism in neurons and astrocytes. Disruptions in glutamate metabolism can precipitate an overabundance of glutamate, which in turn may trigger activity-dependent modulation of glutamate receptors. This phenomenon has been observed in *Drosophila*, where an increase in presynaptic glutamate concentrations leads to a notable reduction in both the number of glutamate receptors and the size of the synaptic field (*41*). Similarly, glutamate treatments have been shown to cause a loss of postsynaptic glutamate receptors (*42*). In mammalian systems, prolonged exposure to glutamate has been demonstrated to specifically reduce RNA editing levels of AMPA receptors in primary cortical neurons (*43*). These findings suggest that a compensatory downregulation mechanism may be responsible for the observed decrease in glutamate receptors in response to excess glutamate.

In our study, we revealed that an overproduction of glutamate resulting from *ubr-1* mutations leads to the downregulation of IVM-targeted GluCls, which is subsequently linked to IVM resistance in *C. elegans*. This contrasts with previously elucidated mechanisms of IVM resistance, such as alterations in IVM-receptor binding sites that diminish their affinity, null mutations in IVM-targeted receptors, the upregulation of cellular efflux pumps, and enhanced drug degradation (*2*). Our research introduces a novel metabolic mechanism involving glutamate receptors. We demonstrated that the levels of IVM-targeted receptors are subject to modulation via the glutamate metabolic pathway. This finding highlights the potential for glutamate accumulation, whether through metabolic dysregulation or exogenous administration, to confer IVM resistance, thereby offering fresh perspectives on the mechanisms behind anthelmintic resistance. Our findings therefore reveal a new UBR-1-mediated pathway via GluCls that connects glutamate metabolism with IVM resistance.

Given the high degree of conservation in glutamate pathways between *C. elegans* and parasitic nematodes, the implications of our findings are far-reaching. This highlights the importance of developing new pharmacological agents aimed at modulating glutamate metabolism for the control of parasitic nematodes. Moreover, altering environmental glutamate concentrations—whether through modifications in the physiological environment of nematodes or changes in the host’s diet—may increase the effectiveness of traditional macrocyclic lactone anthelmintics.

### Expanding parasitic control through UBR-1 and glutamate clearance

Although it remains to be seen what percent of Ivermectin-resistant parasites in the wild have disrupted glutamate homeostasis, there is a need for genetic markers of ivermectin resistance in livestock parasites that can be used to better track resistance and to tailor drug treatment. The discovery of UBR-1 as a resistance gene in *C. elegans* will provide a candidate marker that can be followed up in parasitic control.

Pharmacologically, the recommendation for combination anthelmintic treatments to slow the emergence of anthelmintic resistance is well established, as they cover a similar range of parasites but act via different mechanisms (*2, 44*). Our research proposes an innovative approach for parasite prevention: the clearance of glutamate. The use of ceftriaxone, which has been shown to completely recover IVM sensitivity in *ubr-1* mutants, presents a viable option to mitigate anthelmintic resistance prompted by excess glutamate. A combination treatment that includes ceftriaxone and IVM-like anthelmintics is advisable. Moreover, the concurrent use of ceftriaxone with other antiparasitic drugs may prove effective in treating AR parasites.

This dual-treatment strategy offers several benefits, such as the widespread availability of ceftriaxone, its established FDA approval since 1982 (*45–47*), the relatively low dose needed to increase IVM sensitivity, and its cost-effectiveness, particularly in the developing world.

Although the dynamics of anthelmintic resistance may vary among different hosts, our study contributes a fresh outlook by revealing a new mechanism of IVM resistance and advocating alternative preventative measures. We hope that these insights will attract the attention of global health authorities such as the World Health Organization and spark additional research in the realm of parasite control.

## Materials and methods

### *C. elegans* strains and transgenic lines

*C. elegans* strains were cultured at 22°C on nematode growth medium (NGM) plates seeded with *E. coli* OP50 as a food source (*48*). The wild-type Bristol N2 strain was used, and the *ubr-1(hp684)* strains were sourced from an EMS mutagenesis screen. Additional *ubr-1* alleles were generated via the CRISPR-Cas9 system (*29*). Other genetic mutants were obtained from the *Caenorhabditis Genetics Center* (CGC). Transgenic worms were generated through microinjection via standard protocols, with target DNA plasmids at ∼50 ng/µL coinjected with the marker plasmid P*myo-2*::RFP at 5–10 ng/µL. **Table S1** shows a detailed list of the strains used in this study.

### Ivermectin sensitivity assay

Ivermectin and avermectin (product number: PHR1380/31372) were purchased from Sigma Aldrich (St. Louis, MO, USA) and then diluted in dimethylsulfoxide (DMSO). Doramectin (product number: D127678) was purchased from Aladdin (Shanghai, China). A stock solution of 40 mM IVM was prepared in 1 mL of DMSO. Initial ML resistance assays utilized IVM, avermectin, and doramectin at 5 ng/mL. Standard NGM plates with OP50 and 0.03% DMSO were used, with varying IVM concentrations for tests. L4 hermaphrodites were synchronized and exposed to IVM for viability assays. After 20 hours, the body length, pharyngeal pump rate, and locomotion velocity of the survivors were analyzed via a stereomicroscope (Axio Zoom V16, Zeiss). Each experiment was independently replicated at least three times.

### Animal viability and motor analysis

Animal viability was assessed by calculating the relative residual ratio: the number of viable worms after 20 hours of IVM incubation divided by the initial worm count. Viability (%), body length (mm), and velocity (µm/s) were measured via a custom MATLAB script (*49*).

For movement analysis, IVM-exposed worms were transferred to fresh OP50 plates. For locomotion velocity analysis, a single hermaphrodite was transferred to a 60 mm imaging plate. One minute after the transfer, a three-minute video of the crawled animal was recorded on a modified stereomicroscope (Axio Zoom V16, Zeiss) with a digital camera (acA2500-60um, Basler). Postimaging analyses utilized an in-house written MATLAB script. The central line was used for tracking. Images for velocity analysis from each animal were divided into 33 body segments. The midpoint was used to calculate the body length and velocity between each frame. The pharyngeal pumping frequency was directly calculated manually via video as pumps per minute (min^-1^). Behavioral assays (including pumping frequency) were conducted on standard culture plates with freely moving worms.

### Pharmacological pretreatment

For glutamate application, glutamate (product number: G1251, Sigma Aldrich) was dissolved in NGM at final concentrations of 0.2 mM, 1 mM, 5 mM, 10 mM, 20 mM, and 40 mM. Wild-type and *ubr-1* animals were exposed to glutamate from the L1 to L4 stages and then assessed after 20 hours on glutamate and IVM plates. The same experimental procedure was used for γ-aminobutyric acid (5 mM, product number: A2129, Sigma Aldrich) and aspartate (5 mM, product number: A6202, Macklin, Shanghai, China).

In parallel, *ubr-1* mutants were treated with ceftriaxone sodium (product number: TC6154, Biobomei, Hefei, China) at final concentrations of 0.05 µg/mL, 0.5 µg/mL, 5 µg/mL, 50 µg/mL, and 500 µg/mL. Viability and motor ability were evaluated after L4 worms were transferred to plates containing ceftriaxone and ivermectin (5 ng/mL) for 20 hours.

### Molecular biology and RNA interference

All expression and rescue plasmids were generated via the multisite gateway system (*50*). Three entry clones (Slot1, Slot2, Slot3) corresponding to the promoter, target gene and fluorescent marker genes were recombined into pDEST™ R4-R3 Vector II via the LR reaction, and expression constructs were subsequently obtained. The expression patterns of *ubr-1*, *avr-15*, *avr-14* and *glc-1* were examined via promoter elements of 1.6 kb, 2.7 kb, 3.0 kb and 1.14 kb, respectively. DNA fragments were PCR-amplified from N2 genomic DNA.

Double-stranded RNA (dsRNA) interference technology was used to suppress gene expression in specific tissues. Essential to this process is the expression of dsRNA, which is critical for achieving targeted neuronal or tissue-specific RNAi in *C. elegans* (*51*). The plasmid for RNAi was obtained by constructing Slot2 via the BP reaction of a multisite gateway system. The exon-rich region is usually selected as the interfering region (500-700 bp), the sense strand is selected as the target gene for forward interfering sequences (RNAi, sense), and the antisense strand is selected as the target gene for reverse interfering sequences (RNAi, antisense).

Detailed plasmid and primer information is available in **Tables S2-S3**.

### RNA sequencing analysis

For RNA-seq analysis, synchronized L4 worms were washed with bath solution at least 3 times and harvested via centrifugation. Total RNA was isolated via the freeze-thaw method, and different samples were provided to Novogene for RNA-seq analysis. Transcripts per million (TPM), a standard gene expression normalization method that adjusts the expression of a gene to the number of transcripts per million (*52*), was used to indicate quantitative gene expression. The TPM value takes into account the length of the gene and the sequencing depth and converts the expression of a gene to the number of transcripts per million by dividing the count value of each gene by its length and normalizing appropriately.

### Fluorescence microscopy

L4 stage worms were immobilized with 2.5 mM levamisole (Sigma-Aldrich, USA) on thin layers of agarose for imaging with a laser scanning confocal microscope (Olympus FV3000, Japan) with a 60× oil objective (numerical aperture = 1.35). The intensities of AChR and GABAAR in strains EN208 *kr208* [P*unc-29*::UNC-29::tagRFP] and KP5931 *nuIs283* [P*myo-3*::UNC-49::GFP] were measured. ImageJ software was used to process the images and perform intensity analysis, and the results are reported as the normalized fluorescence intensity per individual and are shown as the mean ± SEM.

### In situ Ca^2+^ imaging

The strain SGA616 [P*myo-2*::GCaMP6s::wCherry] was used to facilitate calcium imaging of the pharyngeal muscles. One-day-old adult hermaphrodites were glued and transferred to bath solution for imaging with a 60x water objective (Nikon, Japan; numerical aperture = 1.0). In the experiment, the epidermis behind head of the worm was gently punctured with a glass electrode (1-3 MΩ) to relieve internal pressure, which helps stabilize the calcium imaging. Serotonin (5-HT, product number: S31021, Yuanye, Shanghai, China) and ivermectin stock solutions were diluted to the desired concentrations via the following bath solution (in mM): NaCl 150; KCl 5; CaCl_2_ 5; MgCl_2_ 1; glucose 10; sucrose 5; and HEPES 15, pH 7.3 with NaOH, ∼330 mOsm. The control bath solution, 5-HT alone, or a combination of 5-HT and ivermectin were administered 30 seconds after initiating the recording via a gravity-driven perfusion system (INBIO MPS-3, China). Fluorescence images were acquired with an excitation wavelength LED at 470 nm and a digital sCMOS camera (Hamamatsu ORCA-Flash 4.0V2) under a wide-field microscope (Nikon LV-TV) at a rate of 10 frames per second for 2 minutes. Data were collected from HCImage (Hamamatsu) and analyzed via Image-Pro Plus 6.0 (Media Cybernetics, Inc., Rockville, MD, USA) and ImageJ (National Institutes of Health). The fluorescence intensity of the region of interest (ROI) was defined as *F*, and the background intensity near the ROI was defined as *F_0_*. The true muscle calcium fluorescence signal was obtained by subtracting the background signal from the ROI. Δ*F/F_0_ = (F - F_0_)/F_0_* was plotted over time as a fluorescence variation curve.

### Statistical analysis

GraphPad Prism 8 was used for statistical analyses and graphing. Dose-response residual ratio (DRR), IC_50_, and EC_50_ values were determined via a sigmoidal Dose-response model. Unpaired Student’s *t* test was used to analyze data between two groups, and ordinary one-way or two-way ANOVA was performed for statistical analysis when multiple groups of data were compared. The p values are indicated as follows: ns, not significant, *p < 0.05, **p < 0.01, ***p < 0.001, ****p < 0.0001. The error bars represent the SEMs.

## Author Contributions

S.G. conceived the experiments and drafted the manuscript. Y.L., L.G., and J.W. conducted the experiments and analyzed the data. W.H. and M.Z. assisted with data analysis and experimentation. M.Z. edited the manuscript.

## Acknowledgments

We thank *Caenorhabditis Genetics Center* and Xia-jing Tong for strains; Ying Wang, Ya Wang, Weicheng Duan and Bo Xiong for their technical support; and Lijun Kang, Xun Huang, and Hong Zhang for their reagents and valuable discussions. This research was supported by the Major International (Regional) Joint Research Project (32020103007 to S.G.), the National Natural Science Foundation of China (32371189 to S.G.), and the National Key Research and Development Program of China (2022YFA1206000 to S.G.).

## Declaration of Interests

The authors declare that they have no conflicts of interest.

---

**Figure S1. Dose- and time-dependent patterns of IVM resistance observed in *ubr-1* mutants.** (A, B) Grid plots depicting the viability of wild-type and *ubr-1* animals treated with different concentrations and durations of IVM. The IVM resistance of *ubr-1* is clearly dose- and time- dependent. The shades of red represent worm viability, with darker shades indicating lower survival rates, based on 100 animals per plate and at least 5 trials. (C) Distribution of locomotion velocity of wild-type and *ubr-1* mutants in the absence and presence of IVM (5 ng/mL).

**Figure S2. IVM resistance across various *ubr-1* mutant alleles.** UBR-1 phylogenetic tree predicting potential functional conservation of *ubr-1* in pathogenic helminthes. (B) Structure of the UBR-1 protein with significant mutants used in this study. Three predicted functional domains (ZnF UBR, RING, UBLC) and amino acid substitutions in *C. elegans* alleles are denoted. (C-F) Quantification of viability, body length, pharyngeal pumps, and locomotion velocity in different *ubr-1* mutant alleles in the presence of IVM (5 ng/mL). All the mutants exhibited varying degrees of IVM resistance. For the viability test in (C), n = 50 animals per plate, which was repeated at least 5 times (5 trials), n ≥ 10 animals in (D-F). ***p < 0.001 by one-way ANOVA. Error bars, SEM.

**Figure S3. Ceftriaxone restores IVM sensitivity in *ubr-1* mutants through mechanisms beyond its antibacterial properties.** (A) OP50 bacteria expressing GFP (OP50::GFP) were used to quantitatively analyze the antibiotic activity of ceftriaxone under the same experimental conditions. *Left*: Schematic diagram of the NGM plate with OP50::GFP. *Right*: OP50::GFP fluorescence intensity under different culture conditions. Control, normal NGM plates with standard culture conditions; +Cef, normal OP50::GFP seeded plate with ceftriaxone (50 μg/mL); Diluted, 15x diluted OP50::GFP seeded plates; Condensed +Cef, 20x condensed OP50::GFP seeded plate with ceftriaxone (50 μg/mL). (B) Quantification of OP50::GFP fluorescence intensity in plates grown under different culture conditions. (C-F) Quantification of all IVM resistance phenotypes under different OP50 levels. *ubr-1*’s IVM resistance is independent of the diluted OP50. Increasing OP50 levels also did not alter the ability of Cef to restore the IVM sensitivity of *ubr-1*. For the viability test in (C), n = 50 animals per plate, which was repeated at least 5 times (5 trials), n ≥ 10 animals in (C-E). Black plots: wild-type, blue plots: *ubr-1* mutants. ns, not significant, **p < 0.01, ****p < 0.0001 by one-way ANOVA. Error bars, SEM.

**Figure S4. Knockdown of UBR-1 induces IVM resistance phenotypes.** (A) Expression of functional UBR-1::GFP, driven by its endogenous promoter, was observed predominantly in the pharynx, head neurons, and body wall muscles with weaker expression detected in vulval muscles and the intestine. Scale bar, 20 µm. (B-E) Double-stranded RNA (dsRNA) interference was used to suppress gene expression in specific tissues (Methods). IVM resistance phenotypes were mimicked by RNAi knockdown of UBR-1 driven by its own promoter (P*ubr-1*), the pharynx muscle-specific promoter (P*myo-2*), the pan-neuronal promoter (P*rgef-1*) or body wall muscle (P*myo-3*). For the viability test in B, n = 50 animals per plate and repeated at least 5 times (5 trials), n ≥ 10 animals in C-E, *p < 0.05, **p < 0.01, ***p < 0.001 by one-way ANOVA. All statistical analyses were performed against *ubr-1* mutant. Error bars, SEM.

**Figure S5. Downregulation of IVM-targeted GluCls in *ubr-1* mutants.** (A) Expression pattern and representative fluorescence intensity of primary IVM-targeted GluCls in wild-type and *ubr-1* animals. AVR-15::GFP was expressed primarily in the pharynx; AVR-14::GFP was expressed in head neurons; and GLC-1::GFP was expressed in the head muscles, pharynx and neurons. (B) Quantitative analysis of the normalized fluorescence intensity of these GluCls in different genetic backgrounds. The intensity of IVM-targeted GluCls was significantly reduced in *ubr-1* mutants. (C) The normalized relative abundance of transcripts of the GluCl genes in wild-type and *ubr-1* animals. (D) Overexpression of AVR-15 reduced the viability of *ubr-1* mutant in IVM (5 ng/mL), thus partially restoring *ubr-1*’s IVM sensitivity. (E) Ceftriaxone (50 μg/mL) treatment rescued the IVM sensitivity in *ubr-1*, but not in *avr-14; avr-15; glc-1* triple mutants. ROI, region of interest; n ≥ 10 animals in each group. ns, not significant, **p < 0.01, ****p < 0.0001 by one-way ANOVA. Error bars, SEM.

**Figure S6. Localization and intensity of AChRs and GABAARs in *ubr-1* mutants.** (A, D) Visualization and fluorescence quantification of the acetylcholine receptor (UNC-29) and GABAA receptor (UNC-49) in both wild-type and *ubr-1* animals. (B-C, E-F) Statistical assessment of the puncta count and mean fluorescence intensity for the respective receptors across various genetic backgrounds revealed no notable differences in the distributions of AChR and GABA_A_R in *ubr-1* mutants. ns, not significant; unpaired Student’s *t* test. The error bars represent the SEM.

**Movie S1. Locomotion and body-length phenotypes of the wild type and *ubr-1* mutants before and after treatment with IVM.** Representative free-moving worms of the wild type and *ubr-1*(*hp684*) mutants with or without IVM (50 ng/ml) treatment for 20 hr. The video shows the worm heading left. Scale bar, 50 µm.

## Notes

### Competing Interest Statement

The authors have declared no competing interest.

### Summary of Updates

Section on Introduction, Results and Discussion updated to clarify; Figure 1, 2, 3, 5 and supplementary Figure S1, S3, S5 are revised.

