## Supplementary figures and images for "UBR-1 deficiency leads to ivermectin resistance in *C. elegans*"

### Figure S1

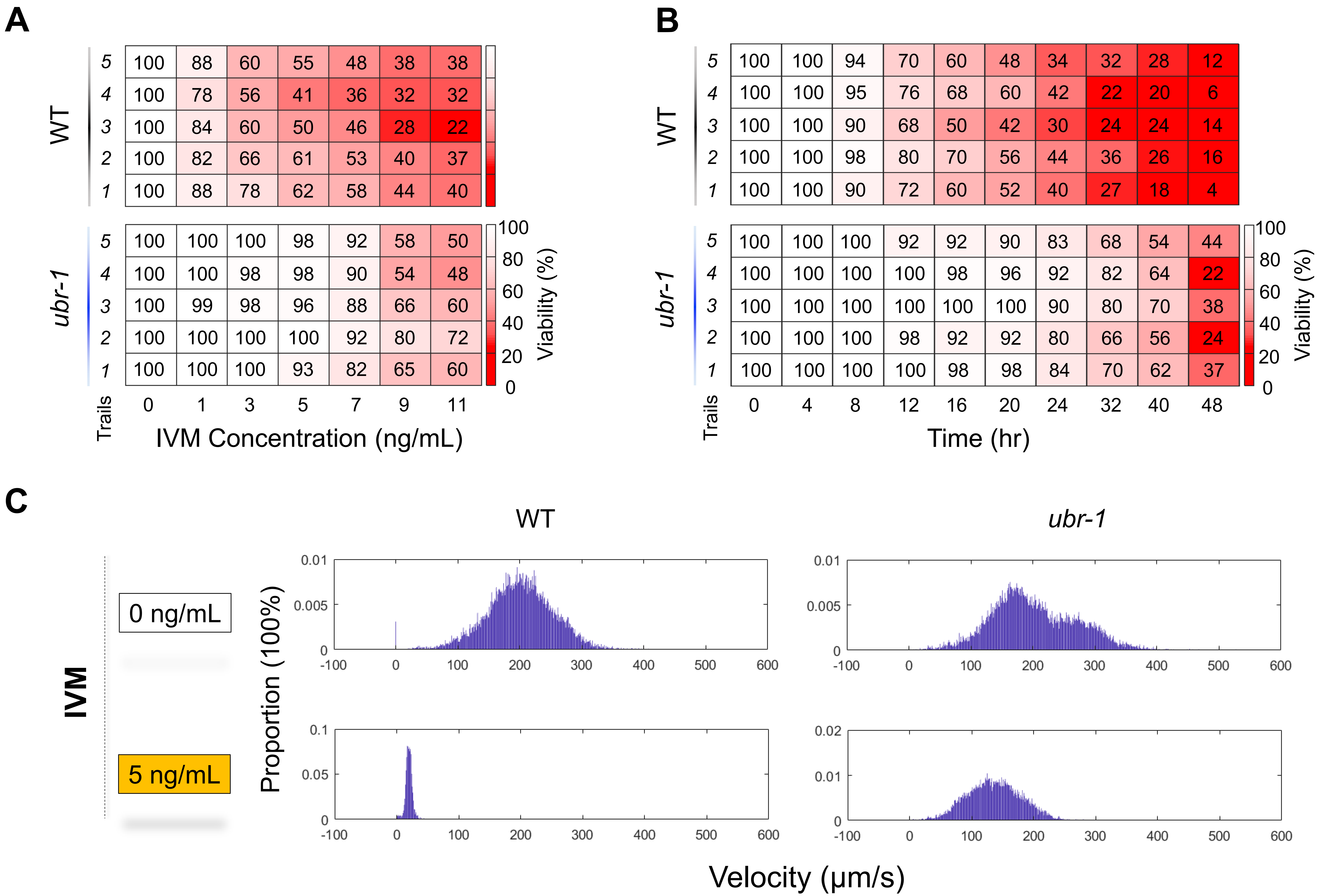

### Figure S2

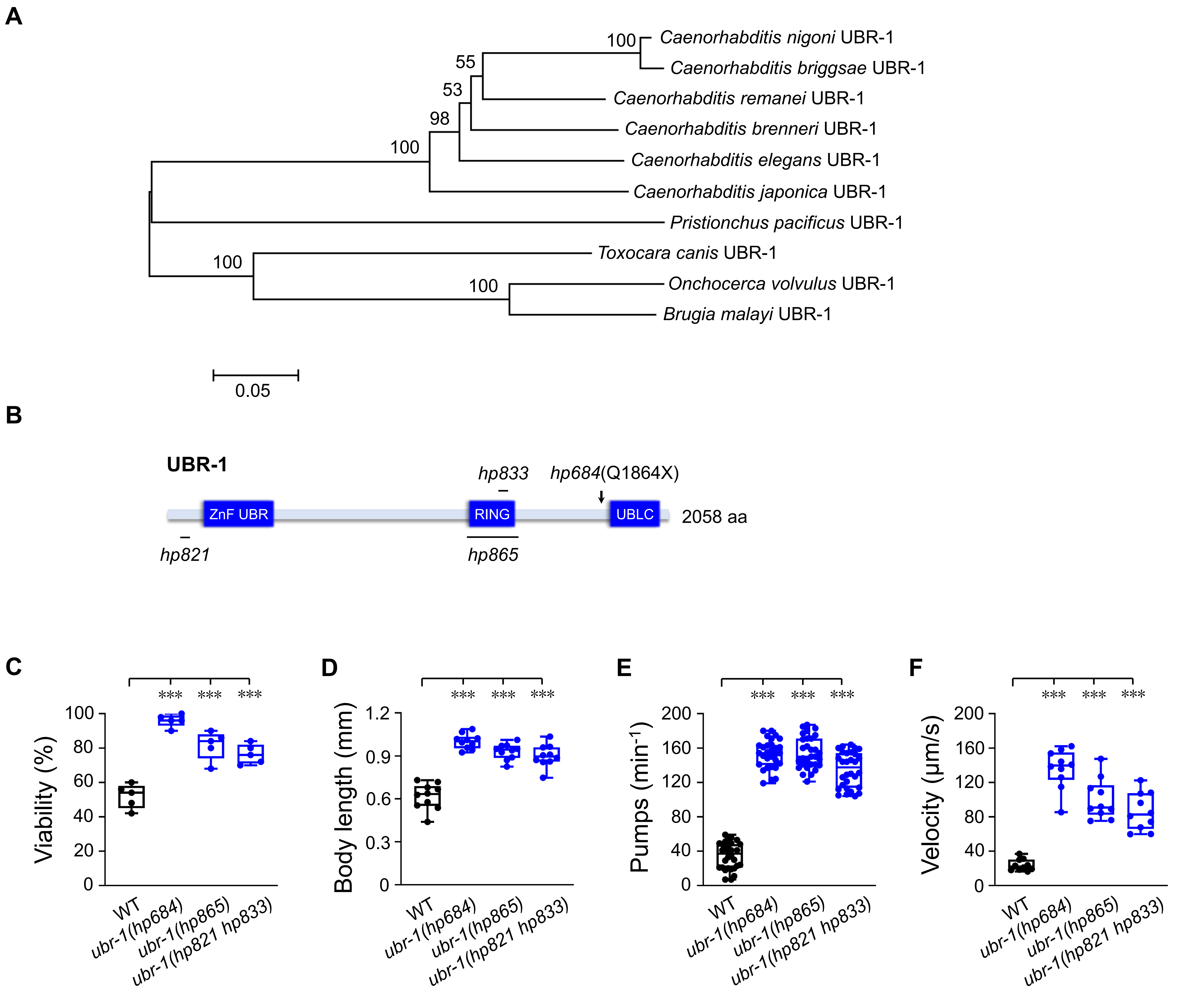

### Figure S3

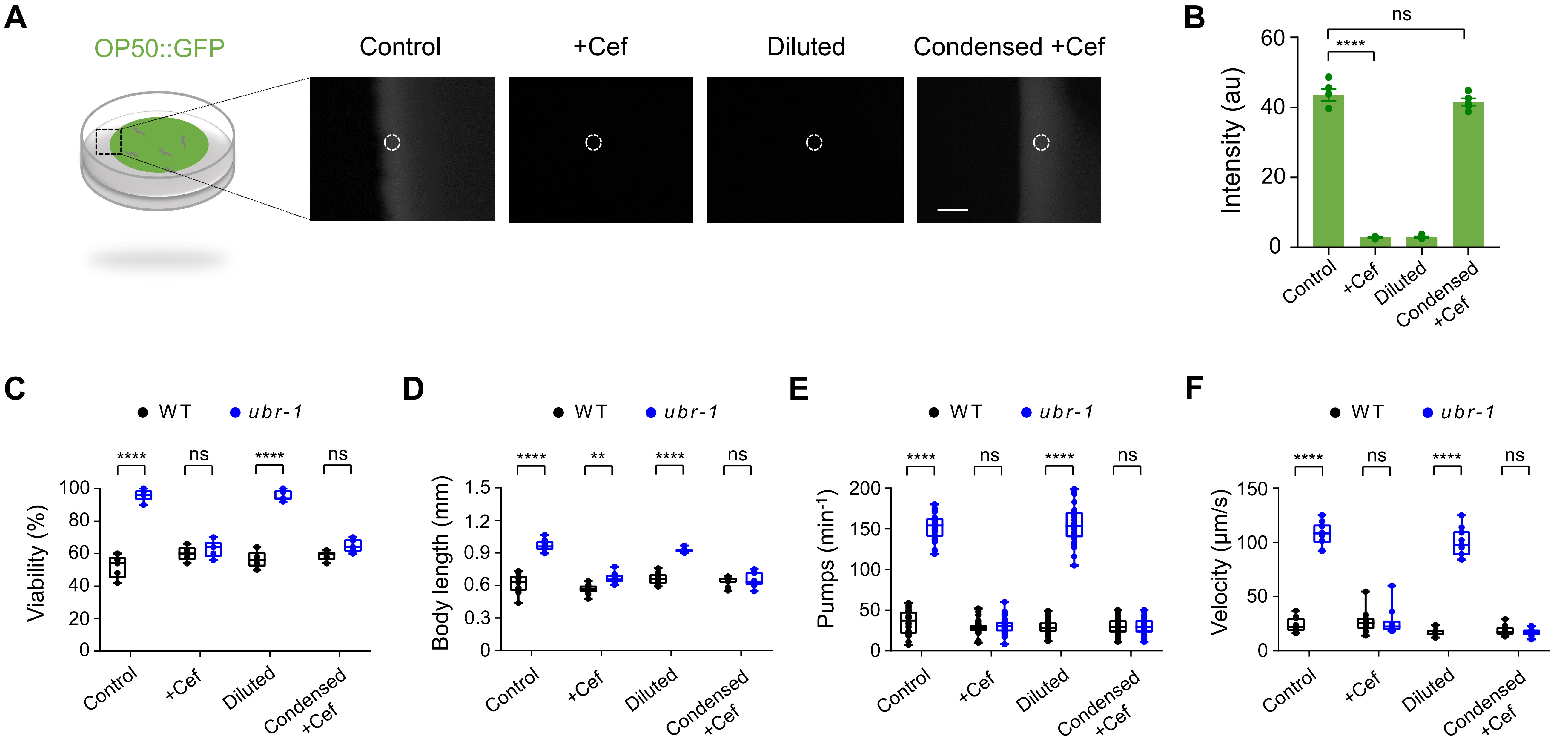

### Figure S4

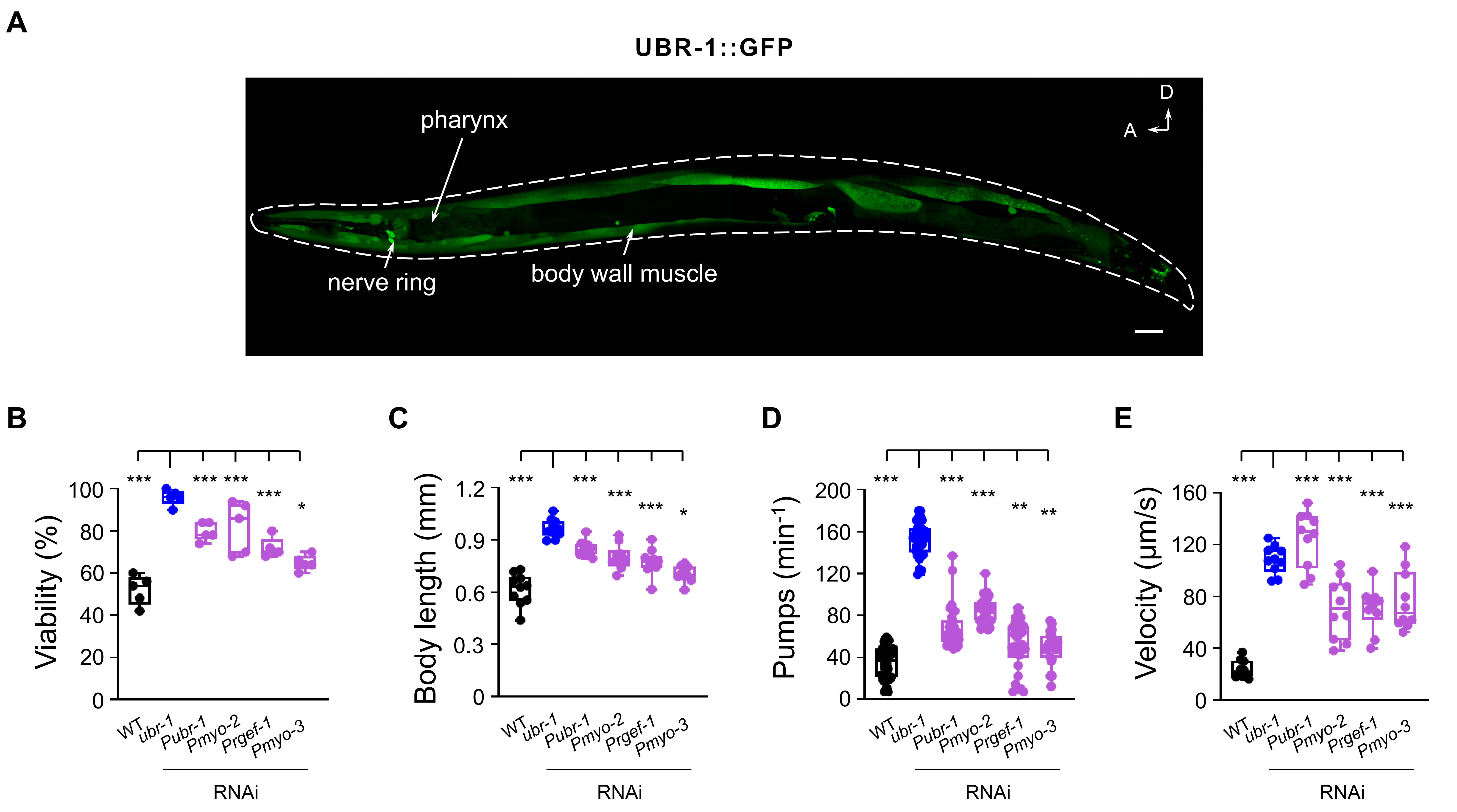

### Figure S5

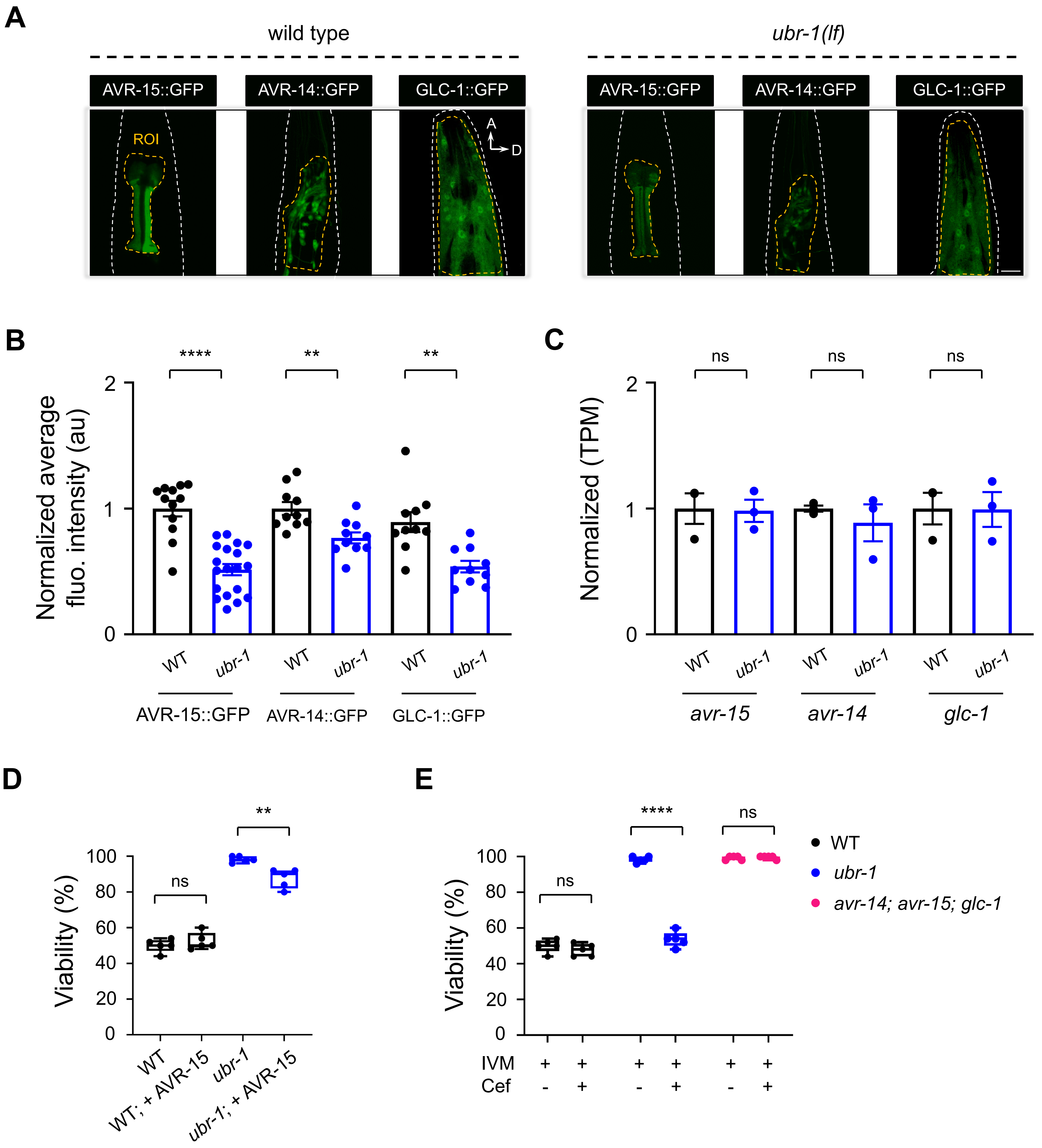

### Figure S6

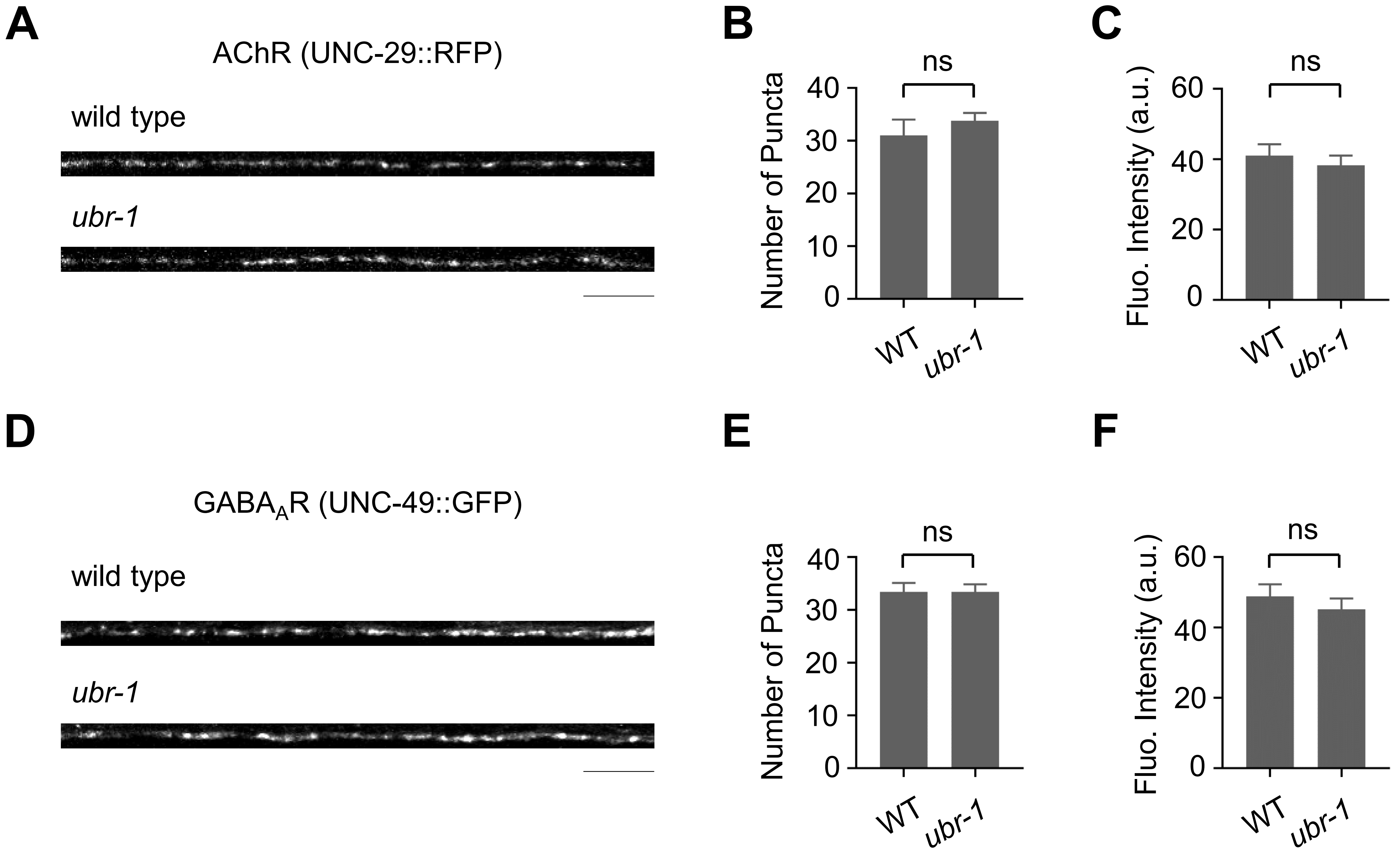
